# Visual activity in primate superior colliculus requires geniculostriate input

**DOI:** 10.64898/2026.04.27.721202

**Authors:** Leor N. Katz, Gongchen Yu, Richard J. Krauzlis

## Abstract

The superior colliculus (SC) is an ancient visual structure whose principle source of visual drive comes directly from the retina. In primates, however, the SC also receives geniculostriate input from primary visual cortex (V1) via the lateral geniculate nucleus (LGN), making it unclear which pathway normally drives visually evoked spiking. Here we tested whether visually evoked spiking in the primate SC depends on retinal signals routed through LGN and V1, rather than on direct retinal input alone. We recorded from macaque SC neurons before and during reversible inactivation of the ipsilateral LGN and found that LGN inactivation nearly abolished visually evoked spiking. This loss was not due to nonspecific suppression of SC, because saccade-related bursts were spared and was observed across SC layers. Magnocellular-biased stimuli, designed to reveal any potential direct retinal drive, failed to produce visually evoked spiking during LGN inactivation. Interhemispheric inhibition (i.e., a “Sprague effect”) was ruled out, because contralateral SC silencing during LGN inactivation did not restore SC visual responses. Consistent with a geniculostriate input route to SC, V1 inactivation also reduced SC visual responses, with effects proportional to the overlap between the V1-induced scotoma and the stimulus representation. Together, these results show that visually evoked neural responses in primate SC depend on retinal signals routed through LGN and V1, and that direct retinotectal input is insufficient to drive SC spiking in the absence of geniculostriate input. These findings revise current models of visual drive to the primate SC and constrain theories of SC-dependent visual behavior.

**Highlights:**

- LGN inactivation nearly eliminates visual but spares movement-related activity in primate SC
- No retinotectal input is detected, even with magnocellular-biased stimuli or in SC input layers
- The elimination of visual activity cannot be explained by interhemispheric inhibition
- Direct retinal projections are not sufficient to drive visual responses in primate SC

## Introduction

The primate superior colliculus (SC) is a midbrain structure that links visual signals to selection and action. It contributes to visual attention ^1^, perceptual decision making ^2^, and orienting movements ^3^, and sits within broader circuits that prioritize behaviorally relevant events. Beyond these established roles, the SC has long been implicated in visual circuits that can guide behavior independently of early visual cortex, both in health and after damage to the primary visual cortex (V1), as in blindsight ^4–7^. Yet, because SC receives visual input through multiple routes, it remains unclear which pathways are necessary to drive visually evoked spiking in SC neurons.

The primary source of visual inputs to the SC in most vertebrate species are direct retinotectal projections. In mice, for example, the vast majority of retinal ganglion cells (RGCs) project to the SC ^8^. In the primate, however, most of the fibers in the optic tract innervate the lateral geniculate nucleus (LGN), which receives inputs from ∼90% of RGCs and conveys these visual signals to V1 via magnocellular and parvocellular channels. V1 in turn provides corticotectal projections to SC that originate predominantly from layer 5 neurons and are strongly driven by magnocellular input ^9,10^. Direct retinotectal projections that are prominent in other vertebrates are also present in primates, but to a lesser extent. Approximately 5–10% of RGCs project directly to primate SC via the retinotectal tract ^11,12^ and this arises largely from magnocellular-type RGCs ^11,13^. Corticotectal and retinotectal inputs prominently target superficial SC and are well positioned to influence the earliest visually evoked responses ^14–16^, whereas additional inputs from extrastriate cortex, frontal areas, basal ganglia, and amygdala shape intermediate and deep layer activity ^2,17^.

In contrast to this well-established anatomy, the functional contribution of retinotectal input to the SC in awake primates is largely unknown. Other than reports in anesthetized animals ^18^, the functional contribution of retinotectal projections in primates has not been demonstrated unambiguously. Despite this lack of evidence, the retinotectal tract is often proposed to provide a fast, V1-bypassing route through which visual stimuli can influence behavior—an idea invoked in accounts of blindsight ^5,19,20^, express saccades ^21–23^, rapid threat processing ^24–26^, and rapid face detection ^6,7,27,28^. In a previous study on face processing in the SC ^29^, LGN inactivation eliminated the visual responses evoked by images of faces and other visual objects. Together, these findings point out the importance of fast visual processing through the SC, but also raise several fundamental mechanistic questions about how SC may draw differentially on geniculostriate versus direct retinal inputs across stimulus regime, SC layers, and visuomotor function.

Our goal here is to define the circuit requirements for visually evoked spiking in the awake primate SC and, more broadly, to reassess what retinotectal projections can contribute in the adult brain. We combine multisite perturbations with targeted analyses to determine how selective the loss of SC visual drive is, how broadly it generalizes across SC layers and stimulus conditions, whether it can be explained by intercollicular inhibition, and whether disrupting early cortical processing by inactivating V1 recapitulates the loss of SC visual responses. Across experiments, LGN inactivation abolishes visually evoked activity almost entirely while sparing saccade-related bursts, including among visuomotor neurons whose visual responses were eliminated. The loss extends into retinorecipient superficial layers, persists across stimuli designed to recruit different retinal ganglion cell populations, and cannot be explained by interhemispheric inhibition because silencing the contralateral SC does not restore visual responsiveness. Finally, we show that the near elimination of visual responsiveness during LGN inactivation could be achieved by inactivating V1, providing direct evidence that early cortical processing is the predominant source of visually evoked spiking in the SC. Together, these results establish a circuit-level account in which robust visual responses in primate SC depend on retinal signals routed through LGN and early visual cortex, placing strong constraints on when and how retinotectal projections may contribute to driving SC activity and informing models of blindsight and subcortical visual circuits.

## Results

We performed a series of experiments to test the proposal that SC visual responses depend on signals routed through early visual cortex. Across 15 experiments performed in two animals, we recorded neuronal activity in the SC using multi-contact recording probes, before and during reversible inactivation of either the LGN, primary visual cortex, or the LGN together with the SC on the opposite side of the brain. By measuring the effects of these causal manipulations on SC activity, we were able to test several alternative hypotheses, as summarized in the sections that follow.

### LGN inactivation eliminates visual but not movement-related activity in SC neurons

LGN inactivation abolishes visually evoked spiking in SC and is thought to indicate a selective loss of visual drive, but it could also reflect a broader, indirect reduction in SC excitability. For example, removing geniculostriate input could shift SC neurons into a hyperpolarized and less excitable state, preventing LGN-independent inputs from driving spiking even if they normally contribute substantially to visual responses. To test whether LGN inactivation rendered SC neurons globally unresponsive or instead selectively affected the visual drive, we compared visually evoked responses with saccade-related activity. If SC neurons were rendered less excitable during LGN inactivation, then movement-related bursts should also be largely eliminated along with the visual responses. Two monkeys either performed an image-viewing task during fixation or freely viewed a nature documentary while making spontaneous saccades. This design allowed us to measure responses to both visual events and saccadic eye movements. SC was recorded at caudal sites with RFs at ∼8° eccentricity, and LGN was injected at sites corresponding to similarly eccentric locations to ensure overlap between the recorded RFs and the region of visual space affected by inactivation, the “LGN-induced scotoma” (Figure 1A).

**Figure 1.**
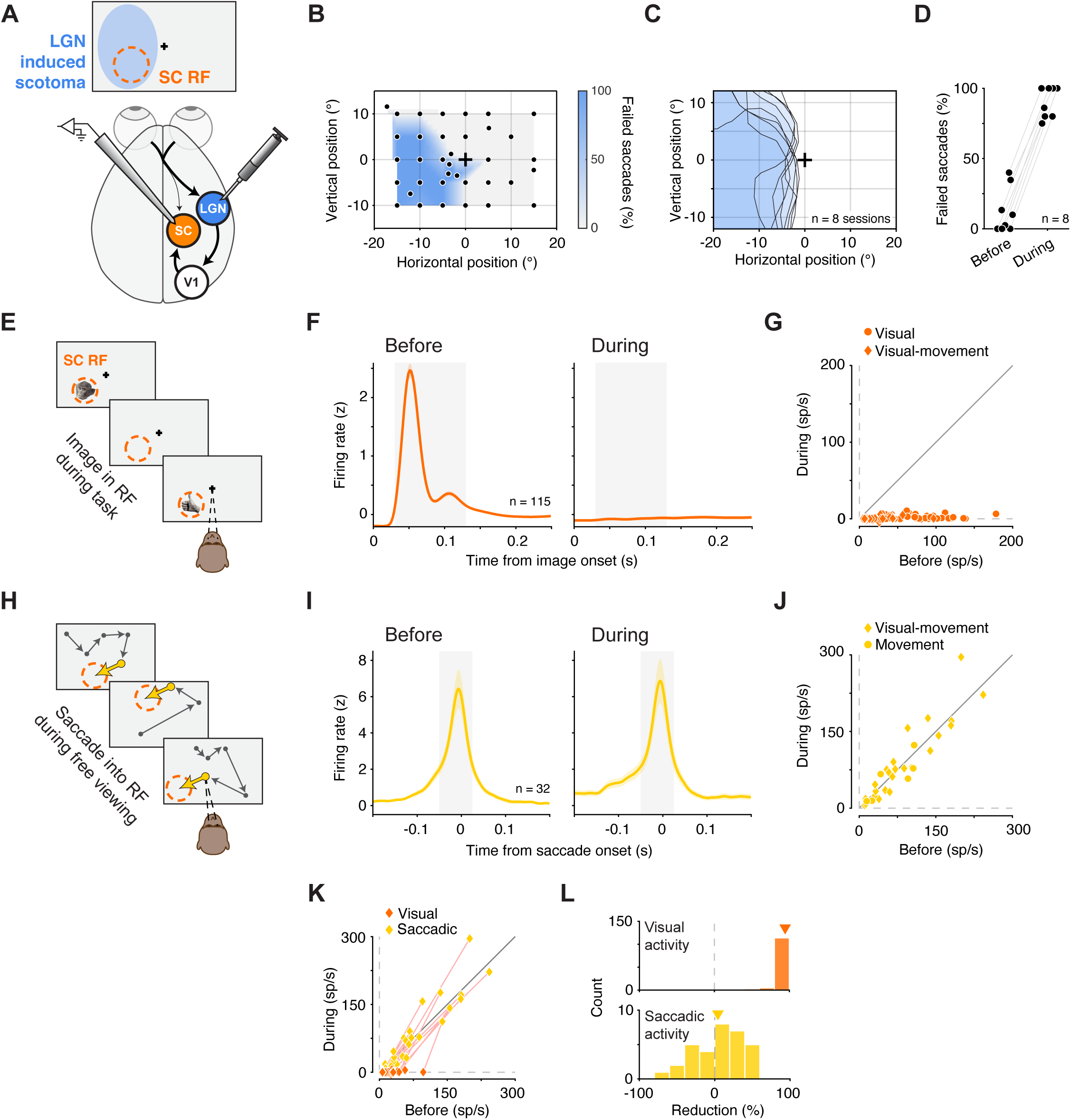
LGN inactivation eliminates visual but not movement-related activity in SC neurons. (A) Experimental schematic. SC neurons were recorded while LGN was reversibly inactivated by muscimol injection at retinotopically matched sites, producing an “LGN-induced scotoma” that overlapped the SC response field (RF). (B) Example session scotoma map measured with a visually guided saccade task. Color denotes fraction of failed saccades to each target location. Cross: fixation location. Individual dots: saccade target locations. (C) Scotomas from 8 LGN inactivation experiments. Shaded contours denote the scotoma measured in each session; cross: fixation location. (D) Percent of failed saccades to targets at the SC RF center before versus during LGN inactivation. Data x-position was jittered for visualization. (E) Image-viewing task used to measure visually evoked activity. Images were sequentially presented in the SC RF during fixation. (F) Population-average normalized firing rate aligned on image onset for visually responsive SC neurons (n = 115) before (left) and during (right) LGN inactivation. Shaded region indicates the visual response window (30–130 ms). (G) Neuron-by-neuron comparison of baseline-subtracted visually evoked responses in the visual response window (panel F) before versus during LGN inactivation. Symbols shapes denote functional class (visual and visual-movement). (H) Free-viewing paradigm used to measure movement-related activity: spontaneous saccades into the RF during viewing of nature documentary videos. (I) Population-average normalized firing rate aligned to saccade onset for movement-responsive neurons (n = 32) before (left) and during (right) LGN inactivation. Shaded region indicates the saccade-response window (−50 to +25 ms). (J) Neuron-by-neuron comparison of baseline-subtracted saccade-related activity in the saccade response window (panel I) before versus during LGN inactivation. Symbol shapes denote functional class. (K) Within-neuron comparison for the visual-movement neurons (n = 25). Neuron-by-neuron baseline-subtracted visual activity (as in G) and of baseline-subtracted saccade-related activity (as in J), before versus during LGN inactivation. Pink lines connect measurements for the same neuron. (L) Distributions of percent reduction during LGN inactivation for visual responses (top; n = 115) and saccade-related responses (bottom; n = 32). Inverted triangles indicate mean. Error bars in (F) and (I) indicate the standard error of the mean (SEM).

LGN inactivation produced a sizable, spatially specific scotoma (i.e., a region of visual space in which monkeys failed to report visual events), consistent with previous reports ^30^. We mapped the scotoma using a visually guided saccade task in which targets tiled the visual field and measured the fraction of failed saccades to each location. In control conditions, failure rates were low; during LGN inactivation, pronounced deficits occurred at specific locations, delineating the scotoma (Figure 1B, example session; Figure 1C, all sessions). We confirmed that the scotoma overlapped the SC RFs by measuring failed saccades to the RF center: before inactivation, 13% of saccades failed on average; during inactivation, failures rose to 90% (Figure 1D, Fisher combined p < 1e-7). Saccades to the ipsilateral hemifield, where no deficit was expected, were unaffected (p > 0.05). With the behavioral impact of LGN inactivation established, we next quantified the effects of LGN inactivation on visual and eye movement-related activity in SC neurons.

Visual activity in SC neurons was measured during image presentation in the RF (Figure 1E). Before LGN inactivation, visually responsive neurons (n = 115) responded robustly to image onset, with an early peak ∼50 ms after onset and a second peak shortly thereafter (Figure 1F, left). During LGN inactivation, visual responses in these same neurons were virtually eliminated (Figure 1F, right). We quantified the evoked visual response (i.e., baseline subtracted) in a 30–130 ms window after image onset (shaded in Figure 1F), which captured the bulk of the visual response. For every neuron in our dataset, visually evoked activity was strongly attenuated during inactivation (Figure 1G; significant reduction in 115 out of 115 neurons with p < 0.05 for each, Wilcoxon rank sum, Bonferroni corrected). On average, visually evoked responses were reduced by 98%, from 43.3 ± 3.2 spikes/s (mean ± 1 SEM) to 0.8 ± 0.2 spikes/s (p < 1e-19, Wilcoxon signed-rank test). This effect held for both functional classes of visually responsive SC neurons—“visual” and “visual-movement” ^31^. For visual neurons, responses were reduced by 98% (from 48.8 ± 3.8 to 1.0 ± 0.2 spikes/s, p < 1e-15). For visual-movement neurons, responses were reduced by 99% (from 22.8 ± 4.0 to 0.4 ± 0.3 spikes/s, p < 1e-5), with no significant difference in percent reduction between classes (p > 0.05, Wilcoxon rank-sum test). Thus, LGN inactivation abolished the visual activity in virtually all visually responsive SC neurons, whether or not the neurons possess movement-related activity.

We further documented the changes in visual activity for the small minority of neurons that retained weak but detectable visual responses during LGN inactivation. Of the 115 visually responsive neurons, 15 (13%) still showed a statistically significant increase in firing after image onset relative to baseline in the 30–130 ms window (p < 0.05, Wilcoxon signed-rank test, Bonferroni-corrected). For these neurons, residual responses during inactivation remained above baseline, but were more than 90% smaller than their pre-inactivation amplitudes (91.8% reduction on average). This residual activity may reflect incomplete suppression of thalamocortical input, a contribution from retinotectal pathways, or some combination of the two. Regardless of the source, it was small and present in very few SC neurons, and observed across both superficial and deep layers of the SC.

In contrast to visual activity, movement-related activity remained intact during LGN inactivation. Movement-related activity was measured during spontaneous saccades into the RF during free viewing (Figure 1H). Before LGN inactivation, movement-responsive neurons (n = 32) displayed sharp motor bursts peaking just before saccade onset and tapering afterward, consistent with classic SC motor activity^32^ (Figure 1I, left). During LGN inactivation, the saccade-related activity of these neurons was mostly unchanged (Figure 1I, right). We quantified saccadic related activity in a baseline subtracted window from 50 ms before to 25 ms after saccade onset (shaded in Figure 1I). Overall, across the population of SC neurons, firing rates did not change significantly between the ‘Before’ and ‘During’ conditions of LGN inactivation (Figure 1J; p > 0.05, Wilcoxon signed-rank test), and the movement-related activity was highly correlated between the Before and During conditions (Pearson r = 0.91, p < 1e-12). Among the 32 individual SC neurons, 6 exhibited a statistically significant reduction, 1 showed a significant increase, and the remaining 25 were not significantly modulated (with p < 0.05, Wilcoxon rank-sum test, Bonferroni corrected). The lack of effect held across functional classes: movement-related activity in neurons of both functional classes, visual-movement and movement, was not significantly altered (p > 0.05 for both, Wilcoxon signed-rank test), and the magnitude of effect did not differ between classes (p > 0.05, Wilcoxon rank-sum test).

The analyses above were based on two, largely non-overlapping, populations of SC neurons — visually responsive neurons (Figure 1E–G) and movement-responsive neurons (Figure 1H–J). This suggests an alternative possibility: that the impact of LGN inactivation might depend on neuron pool rather than on the type of signal reaching the neurons. To test this, we focused on a single pool of SC neurons functionally classified as “visual-movement” (n = 25). Because these neurons exhibit both visual and movement-related activity, this allowed us to perform direct within-neuron comparisons. Plotting visual and saccadic responses before versus during LGN inactivation for visual-movement neurons (Figure 1K) showed that visual activity was nearly abolished (98.6% mean reduction, from 22.8 to 0.3 spikes/s, p < 1e-4, Wilcoxon signed-rank test), whereas saccade-related activity was unchanged (p > 0.05). Thus, the effects of LGN inactivation are specific to visual signals and cannot be explained by neuron pool.

In sum, LGN inactivation nearly abolished visual activity in virtually all visually responsive SC neurons while leaving movement-related activity largely unchanged (Figure 1L). This dissociation, observed even within individual visual-movement neurons, shows that LGN inactivation does not render SC neurons globally unexcitable and that the loss of visual responses cannot be attributed to a nonspecific suppression of SC output. LGN-independent inputs to SC, such as retinotectal projections, should therefore be well positioned to drive residual visual responses if they provide substantial visual drive.

### LGN dependence of SC visual responses is observed across SC depth, including retinorecipient layers

The analyses above demonstrate that LGN inactivation abolished visual responses in SC neurons without rendering SC neurons generally unexcitable, implying that if retinal ganglion cells could drive visual responses in the primate SC, we should have detected it. However, retinotectal projections are not uniformly distributed. They primarily terminate within the upper portion of the superficial layers, especially the upper stratum griseum superficiale just below the stratum zonale ^15,33^. This raises the possibility that residual retinotectal drive might be confined to the most superficial, retinorecipient SC laminae and thus missed by analyses that pool across depth.

We therefore examined the laminar distribution of SC neurons to assess whether any lamina—and especially the retinorecipient superficial layers—retained residual visual activity during LGN inactivation. To evaluate activity across depth, we quantified visually evoked multiunit activity (MUA) on each channel of the 32-channel linear probe, spanning several SC layers. In an example dataset (Figure 2A), before LGN inactivation, visually evoked activity emerged ∼30 ms after image onset and peaked at ∼50 ms on most channels of the probe. In contrast, the dorsal-most channels showed no such activity, as these contacts were deliberately positioned in the quadrigeminal cistern just above the estimated SC surface, ensuring that the channels below sampled the superficial, retinorecipient layers. Visually evoked activity was reduced on the ventral-most channels, consistent with the known functional topography of visual responses over SC depth ^31^. During LGN inactivation, visually evoked activity was abolished on all channels, including the most superficial ones (Figure 2A, right).

**Figure. 2.**
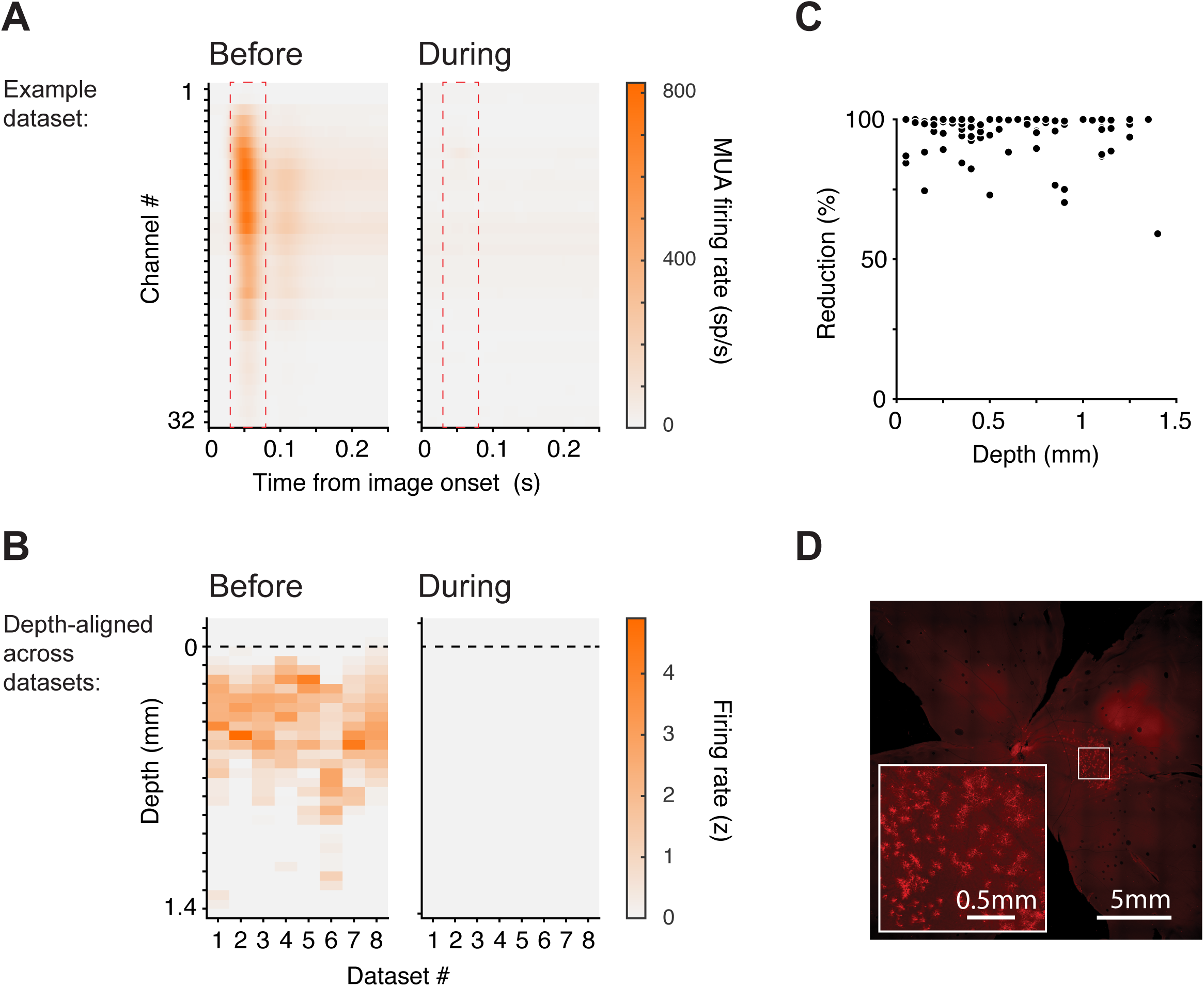
LGN dependence of SC visual responses is observed across SC depth, including retinorecipient layers. (A) Example dataset showing visually evoked multiunit activity (MUA) across the 32 channels of the laminar probe (channel #1 is most dorsal; #32 most ventral), aligned to image onset, before (left) and during (right) LGN inactivation. Dashed red rectangles indicate an early visual response window (30–80 ms). (B) Visually evoked normalized MUA within the early visual response window (panel A), depth-aligned across datasets (see Methods), before (left) and during (right) LGN inactivation. (C) Percent reduction of visually evoked spiking in sorted single units as a function of recording depth (aligned across datasets). (D) Retinal ganglion cell labeling following rAAV2-Retro injection into SC, confirming anatomically intact retinotectal projections. Scale bars as shown.

To determine whether any residual retinotectal drive remained during LGN inactivation, we quantified the evoked response in a window capturing the earliest component of the visual response (30–80 ms after image onset; dashed red rectangles in Figure 2) for each of the eight datasets. We aligned depth profiles to a common reference defined as the first channel with robust visually evoked activity before LGN inactivation (see Methods) and measured visually evoked activity at each channel depth from superficial to intermediate SC layers. Before inactivation, visually evoked activity was typically observed from surface to ∼1 mm below the reference depth, with some variability across datasets (Figure 2B, left). During LGN inactivation, visually evoked MUA was abolished across all probe channels, including the most superficial (Figure 2B, right). To formally test for a depth-localized residual signal, we compared the mean evoked MUA in a superficial band (−50 to 250 µm relative to the reference) to that in a deeper control band (600 to 900 µm) within each dataset and then compared the activity in the superficial and deeper layers using a one-sided signed-rank test. This comparison revealed no evidence for elevated residual activity in superficial channels (p > 0.05). An exploratory sweep across a wide range of plausible superficial and deep band definitions likewise failed to identify any band pair yielding even a trend toward significance. On a neuron-by-neuron basis, we likewise found no evidence for preserved retinotectal input to the most dorsal units: the magnitude of visual response reduction did not covary with recording depth (Figure 2C; r = -0.14, p > 0.05).

A further concern is that the absence of visual responses in retinorecipient SC neurons might reflect damage to these neurons or to retinotectal afferents due to repeated insertions of the recording probes. Anatomical data indicates that this is not the case. One of the animals used in this study later participated in a different study that involved viral vector injection into SC ^34^. In that study, rAAV2-Retro-CBA-Jaws-mCherry-WPRE was injected to test the efficacy of optogenetic manipulation of SC on covert attention. Because this vector is taken up retrogradely at axon terminals, it provides a direct anatomical test for the presence of retinal ganglion cell axons in the SC superficial layers. We found that this injection produced robust labeling in retinal ganglion cells (Figure 2D), confirming that retinotectal projections were structurally intact.

These analyses addressed two concerns about the absence of retinally driven visual responses during LGN inactivation: that we might have undersampled retinorecipient SC neurons and that the direct retinal inputs to SC might have been compromised under our experimental conditions. The result from these analyses demonstrate that the loss of visual responses during LGN inactivation cannot be explained by a failure to sample or maintain retinotectal inputs.

### LGN dependence of SC visual responses persists for motion, flicker, and looming stimuli

Another alternative explanation is that our stimuli may have been poorly suited to engage retinotectal pathways. Retinotectal projections arise from magnocellular ganglion cells ^11,12^, which favor low spatial and high temporal frequencies and motion ^35^, whereas our initial stimulus set consisted of static object and face images. It is therefore possible that the absence of visually driven responses in SC following LGN inactivation reflects insufficient engagement of magnocellular/retinotectal channels.

To test this, we used a set of stimuli explicitly designed to compare parvocellular- and magnocellular-favoring patterns. We first replicated our previous findings with a parvocellular-biased stimulus: a static cluster of black-and-white dots (Figure 3A). As expected, this stimulus elicited strong visual responses in SC neurons shortly after onset, and these visual responses were abolished during LGN inactivation (Figure 3B–C, p < 1e-4, Wilcoxon signed-rank test). We then tested three stimuli designed to favor magnocellular/retinotectal engagement. Two were based on the same black-and-white dot pattern that either moved in unison or flickered (the “motion” and “flicker” stimuli; Figure 3D). A third was a looming pattern (“looming”; Figure 3D), since looming stimuli have been proposed to recruit retinotectal pathways in primates ^36–38^. All three stimuli elicited robust responses in SC neurons before LGN inactivation. During LGN inactivation, however, no visually driven responses were observed for any of the magno-favoring stimuli (Figure 3E–F; motion: p < 1e-4; flicker: p < 1e-4; looming: p < 1e-3; Wilcoxon signed-rank test).

**Figure. 3.**
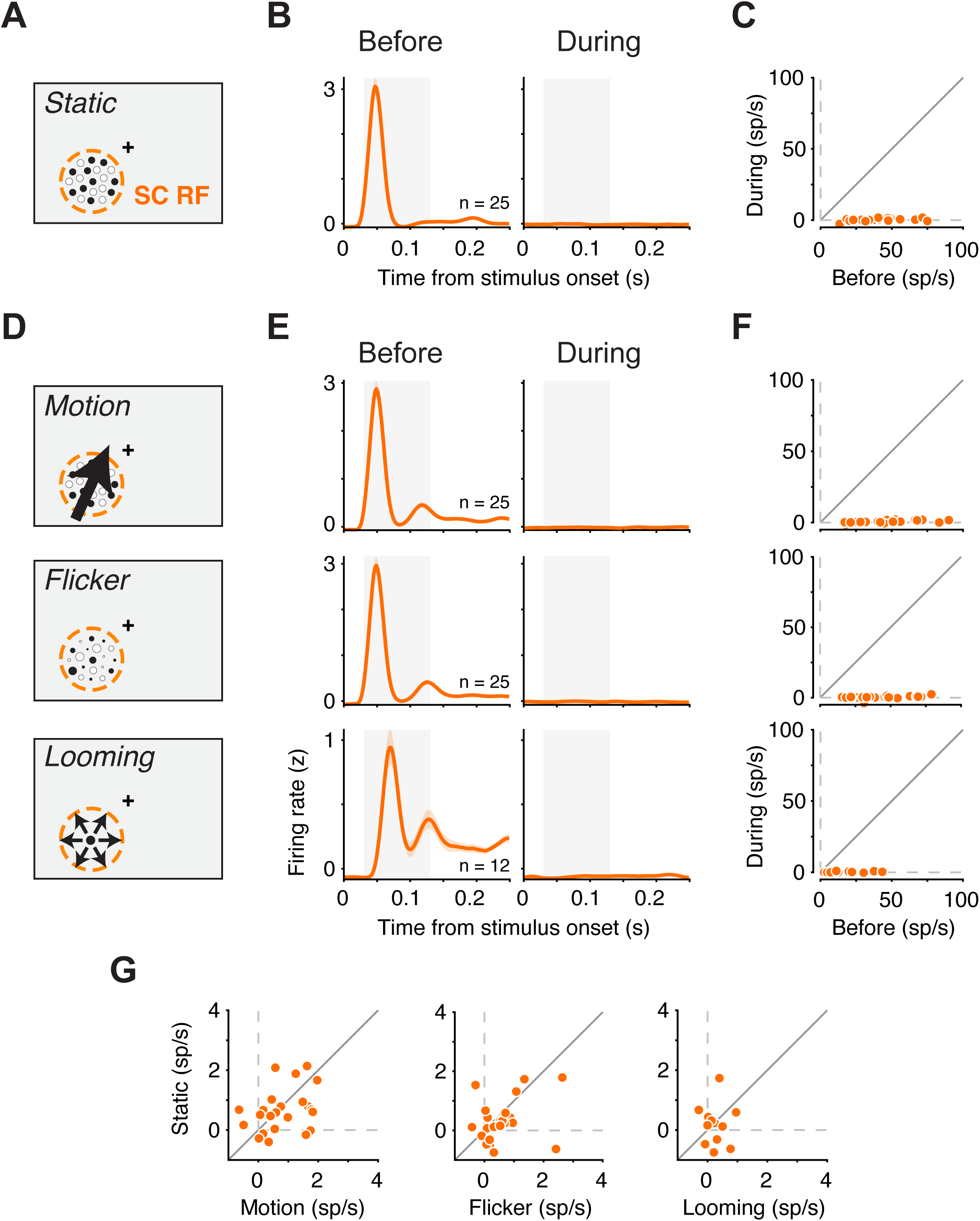
LGN dependence of SC visual responses persists for motion, flicker, and looming stimuli. (A) Parvocellular-biased static dot stimulus used to measure SC visual responses. Cross: fixation. (B) Population-average normalized firing rate aligned to stimulus onset for SC neurons (n = 25) before (left) and during (right) LGN inactivation. Shaded region, visual response window (30-130 ms). (C) Neuron-by-neuron baseline-subtracted activity in the visual response window (panel B) before (left) versus during (right) LGN inactivation. (D) Magnocellular-biased stimuli: motion and flicker versions of the dot pattern, and a looming pattern. (E) Population-average responses for motion, flicker and looming before (left) and during (right) LGN inactivation, aligned to stimulus onset. (F) Neuron-by-neuron responses before versus during LGN inactivation for each stimulus. (G) Residual responses during LGN inactivation for each magnocellular-favoring stimulus versus its parvocellular-biased, static counterpart. Error bars in (B) and (E) indicate SEM.

To test for even subtle relative sparing of magno-favoring signals, we directly compared residual responses during LGN inactivation for each magno-favoring stimulus with its static counterpart. For the motion and flicker conditions, this comparison was especially stringent because the stimuli shared the same dot pattern and differed only in motion or temporal modulation. If retinotectal input selectively supported these magno-favoring signals, residual activity during LGN inactivation should have been larger for motion, flicker, or looming than for the corresponding static stimuli. Instead, residual responses were equally suppressed in all cases, as indicated by the scatter of the data around the unity line in the plot directly comparing visual responses during-inactivation responses for the magno-favoring and static stimuli (Figure 3G; p > 0.05 in all three cases, one-sided Wilcoxon signed-rank test). Thus, even under stimulus conditions tailored to favor magnocellular and looming signals, we found no evidence that retinotectal inputs drive additional spikes in SC during LGN inactivation beyond those elicited by static stimuli.

### Contralateral SC inactivation does not rescue visual responses eliminated by LGN inactivation

Another possible explanation for the loss of visual responsiveness during LGN inactivation is interhemispheric inhibition. Specifically, a Sprague effect-like form of interhemispheric inhibition might be expected to selectively suppress visual responses in the SC ^39,40^. In this framework, inhibition from the contralateral colliculus would be normally balanced by cortical (geniculostriate) drive. However, removing LGN input on one side would tip this balance, allowing the contralateral SC to selectively suppress visual responses in the affected SC—even if retinotectal inputs remain anatomically intact. A key prediction of this model is that inactivating the contralateral SC should relieve this inhibition and restore visual responses in the LGN-inactivated SC.

To test the Sprague-like interhemispheric inhibition account directly, we repeated the LGN inactivation experiments while adding a second inactivation in the contralateral SC (Figure 4A). If the loss of visual activity during LGN inactivation were due to excessive inhibition from the opposite SC, then silencing the contralateral SC should disinhibit the ipsilateral SC and at least partially restore its visual responses. We therefore tested whether the visual activity abolished by LGN inactivation would re-emerge in ipsilateral SC neurons once the contralateral SC was inactivated.

**Figure 4.**
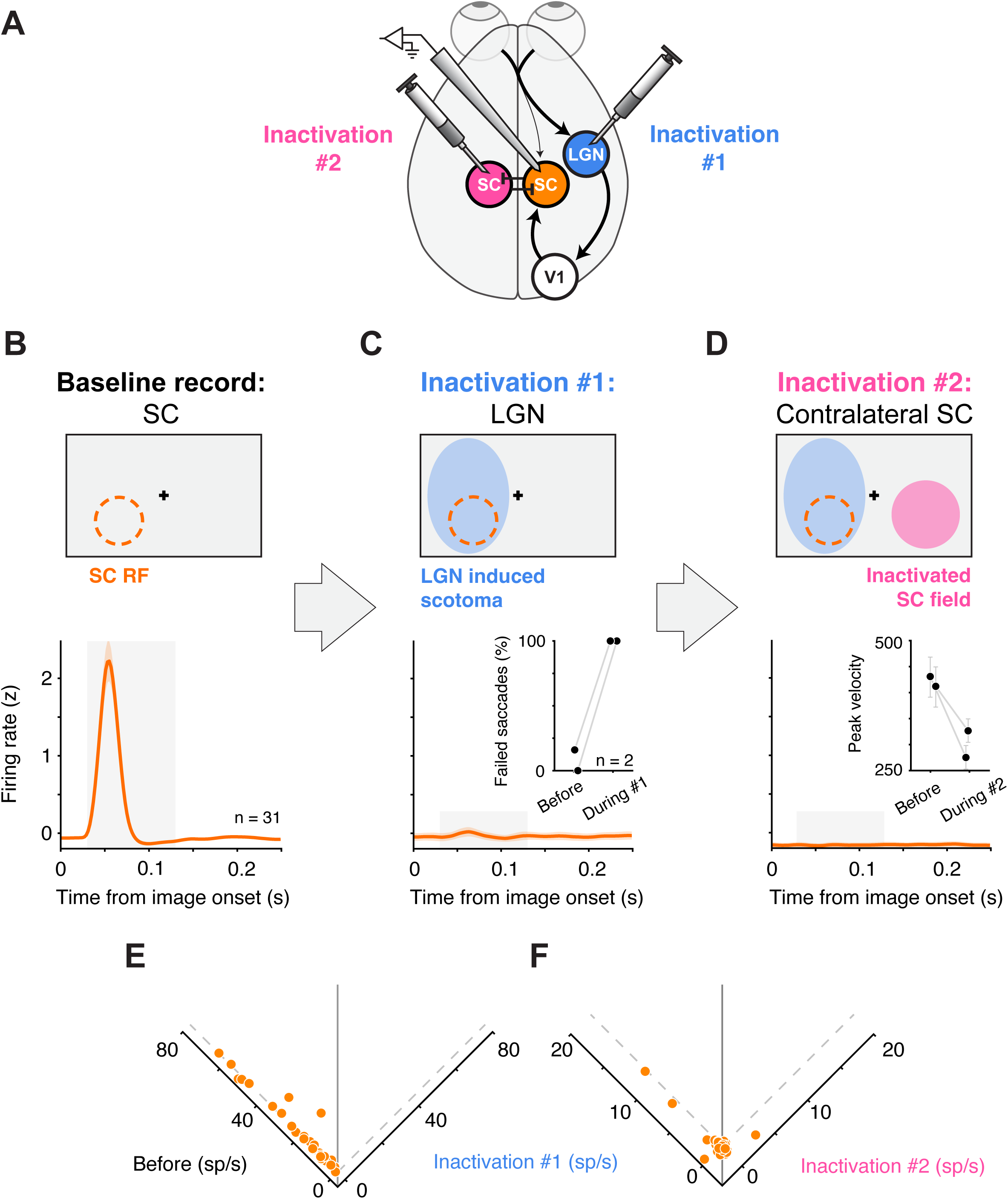
Contralateral SC inactivation does not rescue visual responses eliminated by LGN inactivation. (A) Experimental schematic. SC neurons were recorded while ipsilateral LGN was inactivated (“Inactivation #1”), followed by an additional inactivation of the contralateral SC (“Inactivation #2”). (B) Baseline population response to images presented in the recorded SC RF (n = 31 visually responsive neurons), aligned to image onset. Shaded region, visual response window (30–130 ms). (C) Inactivation #1: LGN inactivation. Top, schematic of LGN-induced scotoma overlapping the recorded SC RF. Inset, percent of failed saccades to targets at the SC RF center before versus during LGN inactivation. Data x-position was jittered for visualization. Bottom, population response to images presented in the RF was abolished during LGN inactivation. (D) Inactivation #2: contralateral SC was inactivated after LGN inactivation. Top, schematic showing the “inactivated SC field” in the contralateral hemifield to the LGN induced scotoma and SC RF. Inset, saccade peak velocity for saccades directed into the inactivated SC field. Bottom, population visual responses in the recorded SC remain absent during Inactivation #2, when both LGN and the contralateral SC are inactivated. (E) Neuron-by-neuron visual response before versus during Inactivation #1 (LGN inactivation), indicating the loss of visual responsiveness during LGN inactivation. (F) Comparison of residual visual activity during Inactivation #1 (LGN inactivation alone) versus Inactivation #2 (during combined LGN + contralateral SC inactivation), indicating no rescue of visual responsiveness. The axis label for Inactivation #1 is shared amongst panels E and F. Error bars in the population response curves (B, C and D, bottom) and in D, inset, indicate SEM.

Before any inactivation, 31 visually responsive SC neurons exhibited short-latency responses to images presented in their RFs (Figure 4B). We then inactivated the ipsilateral LGN and confirmed that the resulting LGN-induced scotoma overlapped the SC RFs (“Inactivation #1”, Figure 4C, top) by measuring increased failure rates for saccades to targets within the RF region of the contralateral hemifield (Figure 4C, bottom, inset). The increase in failure rate was significant in each session (Fisher combined p < 1e-7). As before, visual responses in the same SC neurons were essentially abolished during LGN inactivation in both their mean population response (Figure 4C, bottom) and on a neuron-by-neuron bases (Figure 4E, p < 1e-5, Wilcoxon signed rank), while movement-related responses remained unchanged (p > 0.05).

Next, we inactivated the SC in the opposite hemisphere, contralateral to both the recorded SC and the inactivated LGN (“Inactivation #2”, Figure 4D, top). Based on prior electrophysiological mapping, we selected a site in contralateral SC that matched the recording site in ipsilateral SC, so that the visual field region affected by SC inactivation (the “inactivated SC field”) lay in a roughly mirror-symmetric location relative to the ipsilateral SC RFs. We verified the contralateral SC inactivation using the established method ^41^ of observing a reduction in peak velocities of saccades directed to the center of the inactivated SC field (Figure 4D, bottom, inset); the reductions in peak velocity were significant in both sessions (p < 0.05, Wilcoxon rank-sum). With both the ipsilateral LGN and contralateral SC inactivated, we continued to record responses of ipsilateral SC neurons to images presented in their RFs. Contrary to the Sprague effect prediction, contralateral SC inactivation did not rescue visual responses in the LGN-inactivated SC (Figure 4D, bottom). Visual responses remained absent, and residual visual activity was not larger following SC inactivation than during LGN inactivation alone (Figure 4F, p > 0.05, Wilcoxon signed-rank test). We conclude that an interhemispheric inhibition mechanism does not explain the loss of SC visual responsiveness during LGN inactivation.

### V1 inactivation strongly attenuates visually evoked spiking in SC neurons

The LGN inactivation experiments demonstrate that SC visual responses are abolished when thalamic input is silenced and cannot be explained by residual retinotectal drive. Because LGN has no known direct projection to SC in primates, its influence on SC must be mediated by downstream targets, most prominently V1, which provides dense anatomical inputs to superficial SC ^9,14^. To verify the involvement of V1, we tested whether visual responses in SC were also reduced or eliminated when portions of V1 are inactivated (Figure 5A).

**Figure. 5.**
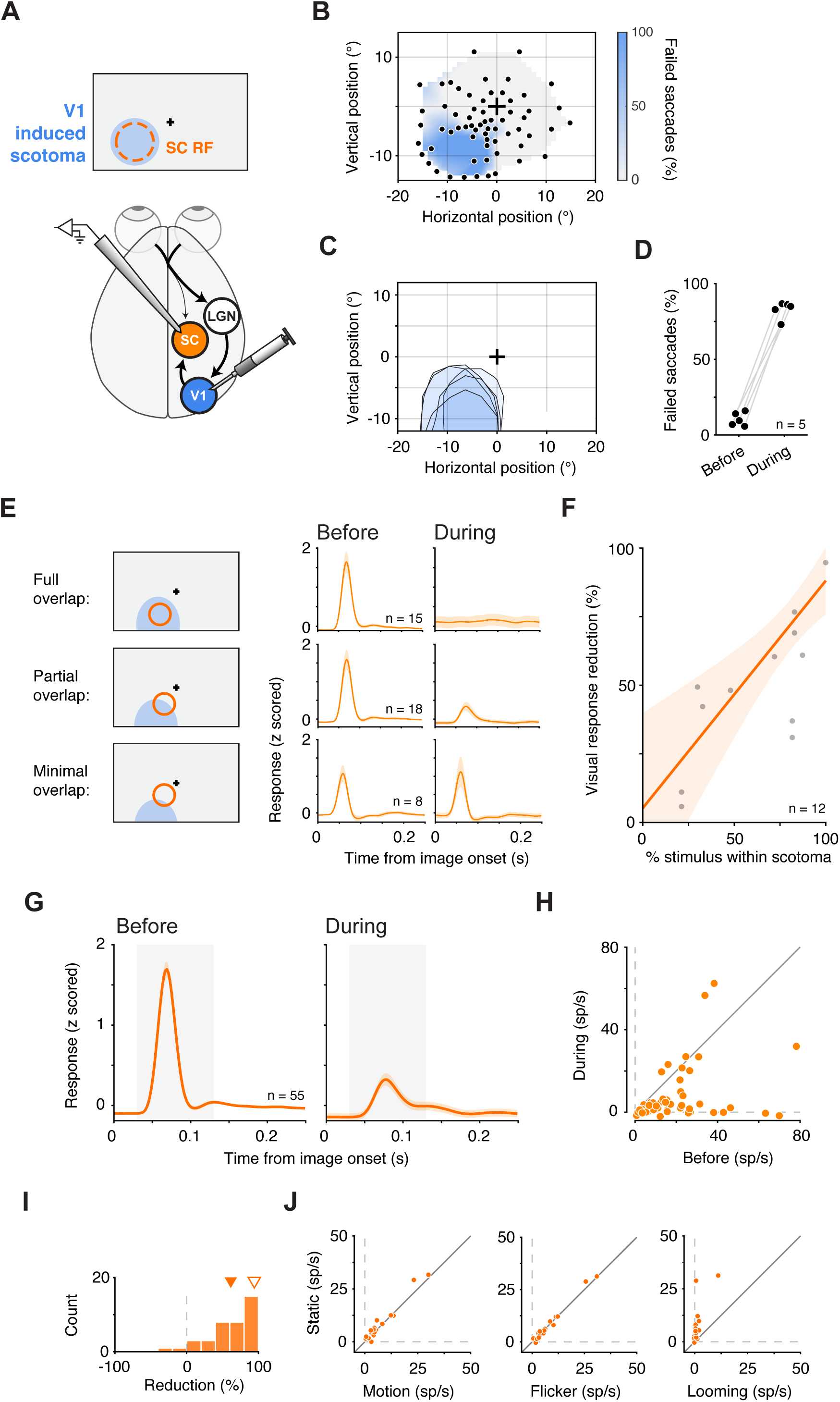
V1 inactivation strongly attenuates visually evoked spiking in SC neurons. (A) Experimental schematic. SC neurons were recorded while V1 was reversibly inactivated at retinotopically matched sites, producing a V1-induced scotoma that overlapped the SC RF. (B) Example session scotoma map measured with a visually guided saccade task; same format as Figure 1B. (C) Scotoma outlines across V1 inactivation sessions. (D) Percent of failed saccades to targets at the SC RF center before versus during V1 inactivation. Data x-position was jittered for visualization. (E) Population-average normalized firing rate aligned to image onset for visually responsive SC neurons recorded in sessions with full, partial or minimal overlap between stimulus and scotoma, before (left) and during V1 inactivation (right). (F) Visual response reduction as a function of the fraction of the stimulus area contained within the V1-induced scotoma within the visual response window, 30 – 130 ms after image onset. Each point shows the mean percent reduction for one stimulus–scotoma overlap condition (n = 12 total conditions across five sessions). Orange line, weighted least-squares linear fit of percent reduction versus percent stimulus overlap; shaded region, 95% confidence band. (G) Population-average normalized firing rate aligned to image onset for visually responsive SC neurons recorded across all sessions (n = 55) before (left) and during (right) V1 inactivation. Shaded region indicates the visual response window (30–130 ms). (H) Neuron-by-neuron comparison of baseline-subtracted visually evoked responses in the visual response window (panel G) before versus during V1 inactivation. (I) Distributions of percent reduction of visually evoked activity in the visual response window (panel G) for V1 inactivation. Inverted solid orange triangle indicates mean. Inversely colored inverted triangle indicates the mean for LGN inactivation, for reference. (J) Residual responses during V1 inactivation for each motion, flicker and looming versus its static stimulus counterpart. Error bars in (E) and (G) indicate SEM.

To confirm that V1 inactivation produced a circumscribed visual scotoma, we used the same visually guided saccade task as in the LGN experiments. Following V1 inactivation, targets presented within the affected region of visual space were frequently missed, whereas targets at other locations remained reliably detected (Figure 5B). Across sessions, V1-induced scotomas occupied the lower visual field and were moderate in size (Figure 5C), substantially smaller than LGN-induced scotomas (compare Figure 5C to Figure 1C), consistent with the larger volume of V1 relative to LGN. Before V1 inactivation, the fraction of failed saccades toward the stimulus site was low (10.4% on average) and increased to 82.7% during inactivation (Figure 5D; Fisher combined p < 1e-37), confirming a localized, contralateral loss of visual sensitivity.

We next examined how the impact of V1 inactivation on SC visual responses depended on the degree of overlap between the visual stimulus and the V1-induced scotoma. The location of the visual stimulus relative to the scotoma was varied in several of the five V1 inactivation sessions, producing 12 distinct stimulus-scotoma overlap conditions. In an example session where the stimulus was entirely contained within the scotoma, visually evoked responses in SC neurons whose RFs lay in that region were virtually eliminated during V1 inactivation (95% reduction, p < 1e-14, Wilcoxon signed-rank test), similar to the effect of LGN inactivation (Figure 5E, top). When the stimulus only partially overlapped the scotoma, SC responses were substantially but not completely reduced (78% reduction, p < 1e-3; Figure 5E, middle). Finally, when stimulus and scotoma overlapped minimally, visually evoked responses in SC were largely preserved (11% reduction, p > 0.05; Figure 5E, bottom). Across the 12 stimulus–scotoma overlap conditions, we found that suppression scaled with overlap. Visual response reduction increased monotonically with the fraction of the stimulus contained within the scotoma (Spearman r = 0.78, p < 1e-2), and a weighted linear regression yielded a consistent positive dependence (Figure 5F). Together, these results show that V1 inactivation suppresses SC visual responses in a spatially specific manner, with the degree of suppression proportional to how much of the stimulus representation was inactivated in the visual cortex.

Next, we summarized the overall impact of V1 inactivation on visually responsive SC neurons, focusing on sessions in which the stimulus overlapped the V1-induced scotoma. Specifically, we included the four sessions with full (1 session) or partial (3 sessions) overlap and excluded the single session with minimal overlap. Across these four sessions, we recorded 55 visually responsive SC neurons whose RFs lay near or within the affected region. Before V1 inactivation, image onsets in the RF elicited robust visual responses (Figure 5G). During V1 inactivation, these responses were strongly attenuated, and on a neuron-by-neuron basis response amplitudes were substantially reduced (Figure 5H; p < 1e-6, Wilcoxon signed-rank test).

Across these SC neurons, V1 inactivation reduced visually evoked responses by 61.5% on average, a large effect but still smaller than the 98% reduction produced by LGN inactivation (Figure 5I). This difference raised the possibility that V1 inactivation spares visual signals that bypass V1 but remain LGN-dependent, for example through extrastriate routes involving MT ^42^. To test this, we presented motion, flicker, and looming stimuli on two sessions (n = 19 neurons), reasoning that such conditions should engage these pathways more strongly than static images (same stimulus set as in Figure 3). However, none of these stimuli evoked stronger SC responses than static stimuli during V1 inactivation (Figure 5J; p > 0.05 for each analysis, one-sided Wilcoxon signed-rank test). We therefore found no evidence that the residual activity was sustained by a V1-bypassing, LGN-dependent visual pathway. Instead, the most likely explanation is that this residual activity reflects the inclusion of sessions with partial and minimal stimulus-scotoma overlap, which we pooled with the full overlap session to increase statistical power. Consistent with this interpretation, when the stimulus was fully contained within the V1-induced scotoma, SC visual responses were nearly abolished (Figure 5E, top), approaching the near-complete loss observed during LGN inactivation.

These results show that the impact of V1 inactivation on SC visual activity scales with the extent to which the stimulus representation is removed from V1. Together with the LGN inactivation experiments, they indicate that SC visual responses are largely, and possibly entirely, driven by thalamocortical pathways routed through V1, with no evidence for an additional source of visual drive when the relevant portion of V1 is silenced.

## Discussion

Our results provide a circuit account of how visual signals drive spiking activity in primate SC. First, inactivating LGN abolished visually evoked responses in SC while sparing movement-related activity across functional class (Figure 1), indicating a selective loss of visual drive rather than a global suppression of SC output. Second, the loss of visually evoked spiking during LGN inactivation was observed across recording depths including the superficial retinorecipient layers of SC (Figure 2), arguing against residual retinotectal drive. Third, the same near-complete loss of visually driven spiking was observed for stimuli designed to favor retinotectal engagement, including motion, flicker, and looming stimuli (Figure 3). Fourth, a Sprague effect-like intercollicular suppression mechanism is unlikely, because inactivating the contralateral SC in addition to the ipsilateral LGN failed to rescue visual responses in SC (Figure 4). Finally, inactivation of V1 reduced SC visual responses in a manner governed by stimulus–scotoma overlap, approaching elimination when overlap was complete (Figure 5). Together, these findings constrain both the source of SC visual drive and the circuit mechanisms that could explain its loss under LGN and V1 perturbations.

### From LGN dependence to circuit mechanism

The present study addresses several key questions raised by our earlier work on face processing in the SC ^29^. In that study, we showed that the short-latency preference for faces – as well as most visually evoked activity in the SC – was largely eliminated during LGN inactivation, indicating that retinotectal inputs alone might be insufficient to drive visual responses. However, those findings did not identify the circuit through which visual signals reach SC, nor did they rule out alternative explanations for the loss of visual activity during LGN inactivation. Because SC receives inhibitory inputs from multiple brain regions, one possibility was that retinotectal inputs remained functional but were ineffective because SC neurons were rendered hyperpolarized or otherwise unresponsive. The present results provide clear and compelling evidence against this interpretation: during LGN inactivation, SC neurons retained robust saccade-related bursts, indicating that they remained excitable and capable of generating high firing rates. In addition, simultaneous inactivation of the contralateral SC failed to restore visual responses, ruling out an intercollicular suppression mechanism.

In our earlier work, we proposed that LGN dependence most plausibly reflects an indirect route through early visual cortex, given the absence of a known direct LGN to SC projection in primates. However, that interpretation remained a circuit hypothesis: LGN inactivation could, in principle, disrupt SC visual responses through multiple downstream relays, without revealing which node is the crucial crux of the circuit. The present results provide that missing causal link. The overlap dependence observed during V1 inactivation indicates that when the relevant portion of V1 is silenced, SC spiking collapses, demonstrating that V1 is required for LGN-dependent signals to influence SC in the affected region of visual space. This constraint limits how much any V1-bypassing pathway can contribute to the responses studied here on the acute timescale and under the stimulus and behavioral conditions tested, while leaving open the possibility that alternative routes contribute under different conditions, response readouts, or after longer-term circuit reorganization.

Having established V1 as a critical node, we can now ask which V1-dependent pathway(s) carry visual drive to SC. One possibility is a predominantly direct corticotectal route from V1 (including layer 5 projections) to superficial SC ^43,44^. In this view, V1 inactivation removes the principal cortical drive to SC, and the dependence on overlap reflects the retinotopic specificity of that drive. A second possibility is convergence, in which V1 provides an enabling input that gates or amplifies signals arriving from other cortical areas that also project to SC, such that SC responses reflect combined V1 and extrastriate influences, with V1 remaining necessary even if not solely responsible for all response components. Distinguishing these models will require targeted perturbations of candidate input areas to SC, or multi-site recordings that identify which cortical signals best predict moment-to-moment variability in SC responses.

V4 is an especially interesting candidate relay for V1-dependent signals to SC. Although V4 projects to SC directly ^14,45^, it is unlikely to provide the earliest volley of visually evoked spiking in SC, given that the earliest SC responses occur at V1-like, not V4-like, latencies ^46,47^ and given the density of direct corticotectal projections from V1 to superficial SC ^43,44^. Instead, V4’s position in the ventral stream makes it well suited to shape the content of SC responses once more elaborated feature combinations and category structure are available ^48,49^.

### The function of primate retinotectal input remains unclear

The absence of visually evoked spiking in SC during LGN inactivation across depth and stimulus conditions places a strong constraint on what the retinotectal input can do on its own in the adult primate. For the vast majority of neurons, visually evoked responses were effectively eliminated, and even when residual responses were detectable in a small subset, they were extremely weak and their source was ambiguous. This result is puzzling. A substantial fraction of retinal output targets SC ^12,13,15^, yet in our hands that input does not appear to provide a strong independent spiking drive when geniculostriate pathway is removed. The most parsimonious interpretation is therefore that whatever retinotectal projections contribute in primates, it is not readily expressed as additive evoked spiking under the conditions tested.

At the same time, the anatomy remains difficult to ignore, and our experiments are not designed to test what the retinotectal input might contribute when the geniculostriate pathway is intact. One possibility is that retinal input acts mainly in a cooperative or permissive manner, shaping how other inputs drive SC rather than driving spikes on its own. For example, retinotectal signals could bias excitability or set timing such that V1 corticotectal or extrastriate inputs are more effective, or could bias selection within SC—for example by modulating competitive interactions across the map—effects that may not appear as a large change in mean evoked firing to isolated stimuli ^1,50,51^. These ideas are speculative, but they have a useful common feature: they predict that retinal input could matter during normal vision, while playing little to no role during acute geniculostriate disruption and passive viewing.

Determining the function of retinotectal inputs might require experiments that isolate retinal input to SC without simultaneously removing cortical drive. Pathway-specific perturbations of retinotectal terminals in SC ^52^, performed while the geniculostriate and cortical pathways remain intact, would provide a direct test of whether retinal input measurably alters response gain, timing, reliability, or selectivity. Critically, this approach would test retinal contributions under conditions where they may be expressed only in the presence of intact cortical drive, rather than as a standalone spiking input revealed by acute thalamic disruption.

Another possibility is that the retinotectal projections are most important for the development, and possibly the maintenance, of the retinotopic map in the primate SC, rather than serving a primary role in visual signal transmission. In many vertebrate species, retinocollicular inputs contribute to the establishment and refinement of topographic maps through molecular guidance and activity-dependent processes, a role that might be orthogonal to driving adult visually evoked spiking ^53,54^.

### Implications for blindsight

Retinotectal projections are also implicated in blindsight—the ability to process visual events following damage to the primary visual cortex, V1 ^5,55^. At first glance, our results argue against a major role for direct retinotectal drive in supporting visually evoked SC spiking under intact conditions, since we found no evidence that retinal input alone can drive visually evoked spiking in SC when V1 is inactivated. However, the present experiments were conducted under acute conditions, whereas primate blindsight and other forms of subcortical vision are thought to reflect circuit reweighting over weeks to months. Pathways that appear functionally silent under acute perturbation could therefore be strengthened or unmasked following permanent lesions. Indeed, recordings from SC in monkeys with chronic V1 lesions demonstrate preserved visually driven activity ^56^, raising the possibility that retinotectal or other subcortical inputs become more influential over time. Related work further shows that residual visual behaviors after V1 lesions can depend critically on SC, since subsequent SC inactivation abolishes recovered visually guided saccades and other blindsight-like responses ^19,57^. This point is important for interpreting the present findings: SC dependence in the chronically lesioned brain does not by itself establish that direct retinotectal input drives visually evoked SC spiking in the intact brain. Rather, it is consistent with the idea that SC becomes a necessary node within a reorganized post-lesion circuit. Longitudinal recordings following V1 lesions could directly test this hypothesis by tracking how visually evoked spiking in SC evolves and evaluate whether recovered signals bear signatures consistent with retinotectal drive versus reorganized cortico-subcortical routes.

Circuit reweighting following V1 damage may also recruit pathways other than the retinotectal tract. One prominent candidate is an LGN-dependent pathway that bypasses V1 while still engaging extrastriate cortex, including direct projections from LGN to MT and other dorsal-stream areas that have been implicated in blindsight ^30,42^. Koniocellular projections may contribute to these alternative thalamocortical routes. Visual signals routed through such pathways could influence behavior either directly or via descending projections to SC, without requiring a direct retinotectal input. The involvement of these pathways would be consistent with our findings that SC responses to magnocellular-biased stimuli do not survive LGN inactivation.

### Reconciling awake and anesthetized measurements of SC visual drive

Our results contrast with those of Schiller et al. ^18^. They found, in anesthetized monkeys, that blocking magnocellular LGN laminae abolished visually driven activity at most topographically corresponding SC sites, yet spared responses in the most superficial retinorecipient zone—consistent with retinotectal drive persisting when geniculostriate or cortical drive is disrupted. However, several methodological differences between our studies stand to reconcile the differences. First, neuronal response properties can differ between anesthetized and awake preparations, which has been observed in SC neurons’ color-related responses ^18,46^. Direct retinal input may be present in both preparations, but whether it is expressed as robust visually evoked spiking or subthreshold and/or modulatory could depend strongly on arousal, ongoing activity elsewhere in the brain, and task engagement. Second, response properties may differ across stimuli. Although we probed retinotectal input with stimuli designed to engage the magnocellular RGCs that constitute the retinotectal tract and still observed no residual responses during LGN inactivation (Figure 3), our stimuli are not identical to those used by Schiller et al. and might have recruited superficial retinorecipient responses differently. Third, Schiller et al. emphasized a very narrow superficial zone; differences in how recording sites are positioned and weighted across that thin layer—particularly with a movable single electrode versus a fixed multichannel probe—could influence whether any residual spiking is readily observed. Taken together, these differences suggest that retinal afferents may be present in both cases but expressed as measurable spiking only under particular states and stimulus/recording regimes, leaving open if and how retinotectal input is translated into visually evoked spikes in primate SC.

## Conclusions

Our findings indicate that in awake primates, visually evoked spiking in SC depends on LGN-driven pathways implemented through V1, placing V1 at the center of the circuit that supplies SC with visual drive. This dependence does not negate a role for retinotectal input but instead sharpens what remains unknown: whether retinal projections primarily modulate SC excitability and timing under intact vision, and whether they become functionally unmasked after chronic loss of cortical pathways. More broadly, this organization suggests that residual, subcortical visual abilities are unlikely to be explained by anatomy alone; they must be understood in terms of how retinal inputs interact with cortical pathways, how those interactions depend on brain state, and how the balance between pathways can be reshaped by injury and experience. By clarifying the circuitry that drives SC visual responses, our work provides a roadmap for identifying the mechanisms that could enable subcortical forms of vision, under normal vision ^6,7^ as well as after cortical disruption and recovery ^4,58^.

## METHODS

### EXPERIMENTAL MODEL AND STUDY PARTICIPANT DETAILS

All experiments were conducted in accordance with NIH guidelines and were approved by the National Eye Institute Animal Care and Use Committee. Experimental procedures complied with the U.S. Public Health Service Policy on the humane care and use of laboratory animals. Investigators were not blinded to experimental conditions during data collection or analysis. The study involved two adult male rhesus macaques (*Macaca mulatta*, aged 13–16 years; 9–12 kg). Each animal had been surgically implanted with a plastic headpost and recording chamber, which allowed stable access to the lateral geniculate nucleus (LGN), superior colliculus (SC) and primary visual cortex (V1) for electrophysiological recordings.

## METHOD DETAILS

### Experimental apparatus

Monkeys were seated in a primate chair (Crist Instruments, Hagerstown, MD) and head-fixed within a darkened booth facing a calibrated VIEWPixx display (VPixx Technologies, 100 Hz refresh rate). Display latency was calibrated using a photodiode, and all event timings were corrected accordingly. Experiments were controlled using a customized version of the PLDAPS framework ^59^. Eye position was sampled at 1000 Hz via an EyeLink 1000 infrared eye tracker (SR Research). Monkeys initiated trials by pressing a vertically oriented joystick (CH Products, HFX-10) mounted to the primate chair.

### Guided Saccade Tasks

Delayed saccade tasks were used for RF mapping, functional classification, and for quantifying inactivation-induced behavioral deficits. At the start of each session, we mapped neuronal RFs using a standard visually guided saccade task and functionally classified neurons using a memory-guided saccade task, assigning units to visual, visual-movement, or movement classes (as in Katz et al. ^31^).

In visually guided saccade trials, monkeys acquired central fixation and a peripheral target appeared 0.75–1 s later and remained visible. After a 1–2 s delay, the fixation point disappeared (go cue), instructing the monkey to saccade to the target. Trials were scored as successful when a saccade was initiated within 1 s of the go cue and the endpoint landed within a 3° square acceptance window centered on the target. Trials were scored as failures if no saccade was initiated within the response window or if the endpoint fell outside the acceptance window. To map inactivation-induced visual deficits for LGN and V1 experiments, targets tiled the visual field, and failure rate was computed separately for each target location to generate a spatial map of saccade failures (scotoma; details below). To verify contralateral SC inactivation, we used the same task to quantify saccade kinematics and measured reductions in peak saccade velocity for targets in the affected portion of the visual field.

### Stimulus Viewing Task

Following RF mapping and functional classification, monkeys performed a stimulus viewing task that consisted of either object images, dots (static, flickering or moving) or looming discs. The individual stimuli are described in the Stimulus Set section. For all stimulus types, trials began with a joystick press, triggering the appearance of a 0.25° white fixation point (48 cd/m²) on a gray background (32.6 cd/m²). Fixation was required within a 1.5° square window (not visible to the subject). After 500 ms of fixation, the background switched to a pink noise texture with identical mean luminance and an RMS contrast of 4.38%. Next, 3–5 stimuli were presented sequentially at a fixed location (5-10° eccentricity), each within a 6° circular aperture. Each stimulus appeared for 400 ms, with 400 ms inter-stimulus intervals. At each stimulus onset and offset, the background noise was refreshed to a new texture. Monkeys received a liquid reward for maintaining successful fixation throughout.

### Stimulus Set

*<u>Object stimuli</u>* included 90 grayscale object images, comprising 30 exemplars each from three categories: faces, hands, and human-made objects, described in detail in Yu et al. ^29^. Object stimuli were presented to both monkeys over 15 inactivation sessions.

*<u>Dot stimuli</u>* could be static, flickering, or moving. Stimuli consisted of black and white square dots (0.25°) presented within the 6° aperture, with 85 dots per stimulus. In the static condition, dot positions were held constant throughout the 400 ms presentation. In the flicker condition, dots were replotted at random locations within the aperture with a dot lifetime of 100 ms (lifetime initialized randomly to distribute replots across time). In the motion condition, dots moved coherently at 15°/s in one of 12 directions with direction jitter (SD = 5°); dot lifetime was 100 ms, after which dots were replotted within the aperture. Dots that reached the edge of the aperture within their lifetime wrapped around. Dot stimuli were presented to one monkey over five inactivation sessions.

*<u>Looming stimuli</u>* consisted of a single pixel that appeared in the center of the stimulus aperture and symmetrically expanded to a 6° diameter disc over the duration of stimulus presentation (radial expansion rate of 7.5°/s). The stimulus color alternated between black and white across trials. Looming stimuli were presented to one monkey over four inactivation sessions.

### Spontaneous saccade detection

Spontaneous saccades were defined as uninstructed saccades that occurred during free-viewing epochs in which monkeys viewed nature documentary videos for 5–8 min. Eye position traces were preprocessed to remove periods in which tracking was lost or gaze fell outside the display boundaries (50° × 36°). Saccades were detected using velocity and acceleration thresholds of 30°/s and 8000°/s^2^ and were required to last at least 12 ms (i.e., ≥12 consecutive samples above threshold). To identify saccades directed into a given neuron’s RF (in-RF saccades), each saccade was parameterized as a polar vector from eye position at saccade onset, yielding direction and amplitude. For each neuron, the RF center estimated during RF mapping was represented as a vector from the fixation position, yielding an RF direction and eccentricity. A spontaneous saccade was classified as in-RF if the saccade endpoint landed near the mapped RF center, defined as a direction that differed from the RF direction by ≤10° (circular difference) and an amplitude between 0.8× and 1.5× the RF eccentricity. Using these criteria, approximately 3% of saccades were classified as in-RF (34/1083 on average before LGN inactivation; 26/810 during). Less stringent criteria increased the number of in-RF saccades but also increased within-group variability; more stringent criteria yielded qualitatively similar results with reduced statistical power.

### Reversible inactivation

While recording from SC, we reversibly inactivated either the ipsilateral LGN, the ipsilateral LGN and the contralateral SC, or ipsilateral V1. Targeting the structures was guided by anatomical MRI and functional response properties. Muscimol (5 mg/ml) was injected via a custom injectrode using a syringe pump (Legato, KD Scientific). Overall, we performed 15 inactivation sessions: eight LGN inactivation sessions (four in monkey #1 and four in monkey #2), two combined LGN and contralateral SC inactivation sessions in monkey #1, and five V1 inactivation sessions in monkey #1.

For sessions with a single inactivation (LGN or V1), we first collected an SC dataset prior to injection (“Before”). We then performed the injection, waited ∼20 min after injection end, and mapped the induced scotoma using a visually guided saccade task ^30^. We subsequently collected an SC dataset during inactivation (“During”) using the same stimulus conditions. After completing the During dataset, we re-mapped the scotoma to confirm that the behavioral deficit persisted through the end of the recording.

For LGN inactivation, 0.8–1.2 µl were infused at 0.1 µl/min to the rostral pole of the LGN, an area of the nucleus that corresponds to the peripheral visual field ^60^, such that the inactivation would not disrupt the monkeys’ ability to fixate and perform the task. Injection location within LGN was confirmed histologically in monkey #2 (see Yu et al. ^29^ for details). For V1 injections, 10–11 µl were infused at 0.25 µl/min over two sites in the operculum separated by 3 mm (∼5 µl per site) to affect the lower visual quadrant.

For both LGN and V1 inactivation, the scotoma was defined as the region of visual space in which the monkey failed to initiate saccades to visual targets; scotoma extent is shown in Figures 1C and 5C (see Yu et al. ^29^ for additional details). Our goal was to induce a scotoma large enough to encompass the stimulus location, which matched the location of the recorded SC neurons’ RFs. This goal was achieved on every LGN inactivation session, during which the overlap between scotoma and stimulus location was complete. This was more challenging in V1 inactivation sessions, when the induced scotomas were smaller and often confined to the lower visual field.

Across V1 inactivation sessions, overlap between the stimulus aperture and the V1-induced scotoma varied. For pooled neuronal analyses summarizing the effect of V1 inactivation on SC visual responses (Figure 5G–J), we included the four sessions with partial or complete overlap and excluded the single session with minimal overlap. The minimal-overlap session was only used for the overlap-dependent example analysis (Figure 5E) and for visualization of the full range of overlap values (Figure 5F).

For sessions with dual inactivation (LGN plus contralateral SC), we first collected a pre-injection SC dataset (“Before”), then injected the ipsilateral LGN. After a ∼20 min post-injection waiting period, we mapped the LGN-induced scotoma and collected an SC dataset during LGN inactivation (“During LGN”). We then injected contralateral SC (0.3–0.4 µl at 0.1 µl/min), targeting intermediate and superficial layers at a retinotopic location chosen to yield an inactivated visual field region approximately the mirror-opposite the recorded SC RFs. Following an additional ∼20 min waiting period, we confirmed successful contralateral SC inactivation by measuring reduced peak saccade velocities for saccades directed into the affected portion of the visual field ^41^. We subsequently collected a second SC dataset (“During LGN + contralateral SC”). In dual-inactivation sessions, we verified that the LGN-induced scotoma persisted after both the During LGN and During LGN + contralateral SC datasets.

### Quantification of the induced scotoma

In addition to mapping the spatial extent of the scotoma (Figure 1C and 5C), we quantified the percent of failed saccades into the circular aperture in which the stimulus was presented, before and during the inactivation (Figure 1D, 2C inset, and 5D). For each session, we quantified saccade failures as binomial counts (failed and successful saccades) in two unpaired phases (Before vs. During). Within each session, we tested for a change in failure probability using a two-sided Fisher’s exact test on the 2×2 contingency table (phase × outcome). To obtain a condition-level inference across sessions, we combined the session-wise p-values using Fisher’s combined probability test ^61^, evaluating the combined statistic against a χ² distribution with 2k degrees of freedom (where k = number of sessions).

To estimate the degree of overlap between scotoma and stimulus (Figure 5E, F), we first parametrized scotoma extent from the visually guided saccade failure map and then quantified its intersection with the stimulus aperture. We constructed a 2D visual-field map by binning target locations on a Cartesian grid and assigning each bin the fraction of failed saccades to targets in that bin. We fit the resulting binned map with a rotated two-dimensional Gaussian plus a constant offset background using least-squares optimization (parameters: amplitude, center x0/y0, σx/σy, rotation angle, constant). We then defined the scotoma boundary as the elliptical contour that contained 95% of the Gaussian’s total integral, determined by the fitted center, widths, and rotation. To quantify overlap, we represented the stimulus as a circular aperture at its known location and size in the same coordinate system and computed the area of intersection between the stimulus aperture and this 95% mass contour. Overlap was expressed as the fraction of stimulus area lying within the scotoma (intersection area / stimulus area) and was computed on every session. Note that on some experimental sessions we collected additional datasets with the stimulus repositioned to either increase or decrease the overlap, netting 12 conditions over the 5 experimental sessions.

### Retinotectal tracing

Monkey 1 was injected with a viral vector rAAV2-Retro-CBA-Jaws-mCherry-WPRE (5.0 × 10¹² vg/mL; Duke Viral Core) into the SC. The virus was taken up at axon terminals and retrogradely transported, resulting in expression of the fluorescent reporter in SC-projecting retinal ganglion cells. Details of the virus injection and general immunohistochemical procedures are described in Katz et al. ^34^. Details of the methods as they pertain specifically to retinal immunostaining and imaging are described in Bohlen & Rudzite et al. ^62^.

### SC Recordings

Neuronal data were acquired using an Omniplex-D system (Plexon Inc.) with 32-channel v-probes with 50 µm spacing. Electrodes were advanced using a motorized microdrive (NAN Instruments) to target superficial and intermediate SC layers, based on functional criteria (i.e., observing neurons with visual and visual-movement activity). Putative deep layer neurons (with movement related activity and no visual responses) were also recorded, but in lower proportion, since we preferentially targeted superficial SC layers. Probes were lowered at an angle of 38° from vertical and allowed to settle for ∼1 h prior to data collection to enhance signal stability. The angle of approach was such that most neuronal RFs overlapped, confirmed using 2D Gaussian RF fits. Task stimuli were placed in a position that maximized the overlap of RFs. Overall, SC data were collected over the 15 inactivation sessions: eight LGN inactivation sessions, two LGN and contralateral SC inactivation sessions, and five V1 inactivation sessions.

### Alignment of recording sessions in depth

To align recording sessions to a common depth origin (Figure 2), we estimated the dorsal surface of SC from the emergence of visually evoked multi-unit activity (MUA). For each session, we computed MUA from threshold crossings on the high-pass filtered signal independently on each of the 32 probe channels using a range of negative thresholds (−3 to −6 SD of the channel’s noise, in 0.5 SD steps). For each threshold value and channel, we quantified visually evoked activity on each trial as the difference in firing rate: ΔFR = FR(30 to 130 ms after stimulus onset) − FR(0 to 100ms before stimulus onset), and tested whether ΔFR was greater than zero across trials (right-tailed Wilcoxon signed-rank test). P-values were FDR-corrected across channels within each session and threshold. For a given threshold, the “estimated surface channel” was defined as the most superficial channel that began a run of three consecutive channels with significant visually evoked activity.

To select a single threshold for depth alignment, we compared the MUA-derived estimated surface channel to an independent single-unit-based estimate of surface (defined as the channel at which the most dorsal isolated single unit was recorded) across sessions, and chose the threshold that maximized the correspondence between these two estimates (Pearson correlation across sessions). This procedure selected a threshold of −5 SD. Using this threshold, we aligned depth profiles across sessions by shifting each session’s channel-wise response vector so that the estimated surface channel was co-linear across recordings. Channels shifted beyond the probe extent were set to NaN for population averaging.

## QUANTIFICATION AND STATISTICAL ANALYSIS

### Electrophysiological analysis

Continuous spike data were sorted offline with Kilosort2 ^63^, manually curated in Phy2, and refined using in-house tools (https://github.com/ElKatz/kilo2Tools). Neurons were excluded if they exhibited poor waveform shape, signal-to-noise ratio lower than 1.8 ^64^, unreasonably low firing rates (<1 sp/s), no clear RF (either visual or movement), or if they were not held continuously throughout the Before and During inactivation datasets.

Our main electrophysiological analyses were performed on visually responsive neurons and included both visual and visual-movement classes of SC neurons, based on responses in the functional classification task ^31^. This inclusion criteria netted 115 neurons during the LGN inactivation sessions, 31 neurons during the LGN and contralateral SC inactivation sessions, and 55 neurons during the V1 inactivation sessions. For analyses of movement-related activity, the same functional classification was applied to include only movement and visual-movement neurons, netting 32, 21, and 33 neurons, for the LGN, LGN+contralateral SC, and V1 inactivations, respectively. Only data from successfully completed trials were included in the analyses.

For population peri-event time histograms (Figures 1F, 1I, 2B, 2C, 2D, 3A, 4B, 4E, 5E, 5G), spike counts were computed in 20 ms windows sliding in 1 ms steps. For visualization only, each neuron’s peri-event time histograms was z-score normalized (mean and standard deviation computed across all time bins pooled across trials and conditions for that neuron). All response quantification on a neuron-by-neuron basis used raw firing rates (spikes/s) rather than z-scored values.

To quantify visually evoked and movement-related activity on a neuron-by-neuron basis, we measured event-related changes in firing rate relative to a pre-event baseline (ΔFR). For visually evoked activity, ΔFR was computed as the mean firing rate in a post-stimulus window (30–130 ms after image onset) minus the mean firing rate in a pre-stimulus baseline window (-100–0 ms relative to image onset). For movement-related activity, ΔFR was computed as the mean firing rate in a peri-saccadic window (-50 to +25 ms relative to saccade onset) minus the mean firing rate in a pre-saccadic baseline window (-300 to -200 ms relative to saccade onset). These visual- or saccade-evoked ΔFR measures are reported in Figures 1G, 1J, 1K, 2B, 3C, 3F, 3G, 4E, 4F, 5F, 5H and 5J.

To quantify the effect of causal manipulations on evoked activity, either visual or saccade-related, we computed percent reduction as (ΔFR_Before_ - ΔFR_During_) / ΔFR_Before_ × 100 (Figures 1L, 2C, and 5I). Analyses were restricted to neurons meeting inclusion criteria for the relevant modality (i.e., classified as possessing either visual or movement properties), ensuring ΔFR_Before_ was nonzero for all percent-change calculations.

Group comparisons used non-parametric tests unless otherwise noted. Confidence intervals were obtained by bootstrap resampling (10,000 resamples).

## Acknowledgments

We thank Carlos Mejias-Aponte, Daniel Yochelson, Nick Nichols, Denise Parker, Hayden Warnock and the NIH Neurophysiology Imaging Facility for technical support. We thank Martin Bohlen and Andra Marija Rudzite for assistance with retinal histology and image processing. We are grateful to David Leopold and Christian Quaia for providing feedback on an earlier version of the manuscript, and to Hendrikje Nienborg, Kara Cover, Divya Subramanian, Grant Folkert, Kerry McAlonan and James Cavanaugh for helpful discussions. This work was supported by the National Eye Institute Intramural Research Program at the National Institutes of Health (ZIA EY000511). The contributions of the NIH author(s) were made as part of their official duties as NIH federal employees, are in compliance with agency policy requirements, and are considered Works of the United States Government. However, the findings and conclusions presented in this paper are those of the author(s) and do not necessarily reflect the views of the NIH or the U.S. Department of Health and Human Services.

## Author contributions

Conceptualization: LNK, RJK

Methodology: LNK, GY

Investigation: LNK

Visualization: LNK

Funding acquisition: RJK

Project administration: RJK

Supervision: RJK

Writing – original draft: LNK

Writing – review & editing: LNK, GY, RJK

## References

1. Krauzlis, R.J., Lovejoy, L.P., and Zénon, A. (2013). Superior Colliculus and Visual Spatial Attention. Annual Review of Neuroscience 36, 165–182. 10.1146/annurev-neuro-062012-170249.

2. Basso, M.A., and May, P.J. (2017). Circuits for Action and Cognition: A View from the Superior Colliculus. Annual review of vision science 3, 197–226. 10.1146/annurev-vision-102016-061234.

3. Gandhi, N.J., and Katnani, H.A. (2011). Motor functions of the superior colliculus. Annual Review of Neuroscience 34, 205–231. 10.1146/annurev-neuro-061010-113728.

4. Cowey, A. (2010). The blindsight saga. Experimental Brain Research 200, 3–24. 10.1007/s00221-009-1914-2.

5. Isa, T., and Yoshida, M. (2021). Neural Mechanism of Blindsight in a Macaque Model. Neuroscience 469, 138–161. 10.1016/j.neuroscience.2021.06.022.

6. Johnson, M.H. (2005). Subcortical face processing. Nature Reviews Neuroscience 6, 766–774.

7. Tamietto, M., and De Gelder, B. (2010). Neural bases of the non-conscious perception of emotional signals. Nature Reviews Neuroscience 11, 697–709.

8. Ellis, E.M., Gauvain, G., Sivyer, B., and Murphy, G.J. (2016). Shared and distinct retinal input to the mouse superior colliculus and dorsal lateral geniculate nucleus. J Neurophysiol 116, 602–610. 10.1152/jn.00227.2016.

9. Fries, W. (1984). Cortical projections to the superior colliculus in the macaque monkey: A retrograde study using horseradish peroxidase. Journal of Comparative Neurology 230, 55–76. 10.1002/cne.902300106.

10. Lock, T.M., Baizer, J.S., and Bender, D.B. (2003). Distribution of corticotectal cells in macaque. Experimental brain research 151, 455–470.

11. Perry, V.H., and Cowey, A. (1984). Retinal ganglion cells that project to the superior colliculus and pretectum in the macaque monkey. Neuroscience 12, 1125–1137. 10.1016/0306-4522(84)90007-1.

12. Zheng, Y.J., Adams, D.L., Gentry, T.N., Dilbeck, M.D., Economides, J.R., and Horton, J.C. (2024). Retinal Input to Macaque Superior Colliculus Derives from Branching Axons Projecting to the Lateral Geniculate Nucleus. Journal of Neuroscience 44.

13. Schiller, P.H., and Malpeli, J.G. (1977). Properties and tectal projections of monkey retinal ganglion cells. Journal of Neurophysiology 40, 428–445. 10.1152/jn.1977.40.2.428.

14. Cerkevich, C.M., Lyon, D.C., Balaram, P., and Kaas, J.H. (2014). Distribution of cortical neurons projecting to the superior colliculus in macaque monkeys. Eye Brain 6, 121–137. 10.2147/EB.S53613.

15. May, P.J. (2006). The mammalian superior colliculus: laminar structure and connections. Progress in Brain Research 151, 321–378. 10.1016/S0079-6123(05)51011-2.

16. Pollack, J.G., and Hickey, T.L. (1979). The distribution of retino-collicular axon terminals in rhesus monkey. J of Comparative Neurology 185, 587–602. 10.1002/cne.901850402.

17. Cusick, C.G. (1988). Anatomical organization of the superior colliculus in monkeys: corticotectal pathways for visual and visuomotor functions. Progress in Brain Research 75, 1–15.

18. Schiller, P.H., Malpeli, J.G., and Schein, S.J. (1979). Composition of geniculostriate input ot superior colliculus of the rhesus monkey. Journal of Neurophysiology 42, 1124–1133. 10.1152/jn.1979.42.4.1124.

19. Kato, R., Takaura, K., Ikeda, T., Yoshida, M., and Isa, T. (2011). Contribution of the retino-tectal pathway to visually guided saccades after lesion of the primary visual cortex in monkeys. Eur J Neurosci 33, 1952–1960. 10.1111/j.1460-9568.2011.07729.x.

20. Leh, S.E., Ptito, A., Schönwiesner, M., Chakravarty, M.M., and Mullen, K.T. (2010). Blindsight mediated by an S-cone-independent collicular pathway: an fMRI study in hemispherectomized subjects. Journal of Cognitive Neuroscience 22, 670–682.

21. Anderson, A.J., and Carpenter, R.H.S. (2008). The effect of stimuli that isolate S-cones on early saccades and the gap effect. Proc. R. Soc. B. 275, 335–344. 10.1098/rspb.2007.1394.

22. Koller, K., and Rafal, R.D. (2019). Saccade latency bias toward temporal hemifield: Evidence for role of retinotectal tract in mediating reflexive saccades. Neuropsychologia 128, 276–281.

23. Marino, R.A., Levy, R., and Munoz, D.P. (2015). Linking express saccade occurance to stimulus properties and sensorimotor integration in the superior colliculus. Journal of Neurophysiology 114, 879–892. 10.1152/jn.00047.2015.

24. Inagaki, M., Inoue, K., Tanabe, S., Kimura, K., Takada, M., and Fujita, I. (2023). Rapid processing of threatening faces in the amygdala of nonhuman primates: subcortical inputs and dual roles. Cerebral Cortex 33, 895–915.

25. Liddell, B.J., Brown, K.J., Kemp, A.H., Barton, M.J., Das, P., Peduto, A., Gordon, E., and Williams, L.M. (2005). A direct brainstem–amygdala–cortical ‘alarm’system for subliminal signals of fear. Neuroimage 24, 235–243.

26. Morris, J.S., Öhman, A., and Dolan, R.J. (1999). A subcortical pathway to the right amygdala mediating “unseen” fear. Proc. Natl. Acad. Sci. U.S.A. 96, 1680–1685. 10.1073/pnas.96.4.1680.

27. Le, Q.V., Le, Q.V., Nishimaru, H., Matsumoto, J., Takamura, Y., Hori, E., Maior, R.S., Tomaz, C., Ono, T., and Nishijo, H. (2020). A prototypical template for rapid face detection is embedded in the monkey superior colliculus. Frontiers in systems neuroscience 14, 5.

28. Nguyen, M.N., Matsumoto, J., Hori, E., Maior, R.S., Tomaz, C., Tran, A.H., Ono, T., and Nishijo, H. (2014). Neuronal responses to face-like and facial stimuli in the monkey superior colliculus. Frontiers in behavioral neuroscience 8, 85.

29. Yu, G., Katz, L.N., Quaia, C., Messinger, A., and Krauzlis, R.J. (2024). Short-latency preference for faces in primate superior colliculus depends on visual cortex. Neuron 112, 2814–2822.

30. Schmid, M.C., Mrowka, S.W., Turchi, J., Saunders, R.C., Wilke, M., Peters, A.J., Ye, F.Q., and Leopold, D.A. (2010). Blindsight depends on the lateral geniculate nucleus. Nature 466, 373–377.

31. Katz, L.N., Yu, G., Herman, J.P., and Krauzlis, R.J. (2023). Correlated variability in primate superior colliculus depends on functional class. Communications Biology 6, 540.

32. Wurtz, R.H., and Goldberg, M.E. (1972). Activity of superior colliculus in behaving monkey. 3. Cells discharging before eye movements. Journal of Neurophysiology 35, 575–586. 10.1152/jn.1972.35.4.575.

33. Wurtz, R.H., and Albano, J.E. (1980). Visual-motor function of the primate superior colliculus. Annual Review of Neuroscience 3, 189–226. 10.1146/annurev.ne.03.030180.001201.

34. Katz, L.N., Bohlen, M.O., Yu, G., Mejias-Aponte, C., Sommer, M.A., and Krauzlis, R.J. (2025). Optogenetic Manipulation of Covert Attention in the Nonhuman Primate. Journal of Cognitive Neuroscience 37, 266–285.

35. Shapley, R., and Perry, V.H. (1986). Cat and monkey retinal ganglion cells and their visual functional roles. Trends in Neurosciences 9, 229–235.

36. Billington, J., Wilkie, R.M., Field, D.T., and Wann, J.P. (2011). Neural processing of imminent collision in humans. Proc Biol Sci 278, 1476–1481. 10.1098/rspb.2010.1895.

37. Cléry, J.C., Schaeffer, D.J., Hori, Y., Gilbert, K.M., Hayrynen, L.K., Gati, J.S., Menon, R.S., and Everling, S. (2020). Looming and receding visual networks in awake marmosets investigated with fMRI. NeuroImage 215, 116815. 10.1016/j.neuroimage.2020.116815.

38. Guo, F., Zou, J., Wang, Y., Fang, B., Zhou, H., Wang, D., He, S., and Zhang, P. (2024). Human subcortical pathways automatically detect collision trajectory without attention and awareness. PLoS Biol 22, e3002375. 10.1371/journal.pbio.3002375.

39. Mekhaiel, D.Y., Corneil, B.D., and Goodale, M. (2024). Does face detection in the superior colliculus rely on input from the primary visual cortex?

40. Sprague, J.M. (1966). Interaction of cortex and superior colliculus in mediation of visually guided behavior in the cat. Science 153, 1544–1547. 10.1126/science.153.3743.1544.

41. Lovejoy, L.P., and Krauzlis, R.J. (2010). Inactivation of primate superior colliculus impairs covert selection of signals for perceptual judgments. Nature neuroscience 13, 261–266.

42. Sincich, L.C., Park, K.F., Wohlgemuth, M.J., and Horton, J.C. (2004). Bypassing V1: a direct geniculate input to area MT. Nature neuroscience 7, 1123–1128.

43. Distel, H., and Fries, W. (1982). Contralateral cortical projections to the superior colliculus in the macaque monkey. Exp Brain Res 48, 157–162. 10.1007/BF00237210.

44. Lund, J.S., Lund, R.D., Hendrickson, A.E., Bunt, A.H., and Fuchs, A.F. (1975). The origin of efferent pathways from the primary visual cortex, area 17, of the macaque monkey as shown by retrograde transport of horseradish peroxidase. J Comp Neurol 164, 287–303. 10.1002/cne.901640303.

45. Gattass, R., Galkin, T.W., Desimone, R., and Ungerleider, L.G. (2014). Subcortical connections of area V4 in the macaque. J Comp Neurol 522, 1941–1965. 10.1002/cne.23513.

46. White, B.J., Boehnke, S.E., Marino, R.A., Itti, L., and Munoz, D.P. (2009). Color-Related Signals in the Primate Superior Colliculus. J. Neurosci. 29, 12159–12166. 10.1523/JNEUROSCI.1986-09.2009.

47. Zamarashkina, P., Popovkina, D.V., and Pasupathy, A. (2020). Timing of response onset and offset in macaque V4: stimulus and task dependence. J Neurophysiol 123, 2311–2325. 10.1152/jn.00586.2019.

48. Pasupathy, A., Popovkina, D.V., and Kim, T. (2020). Visual Functions of Primate Area V4. Annual Review of Vision Science 6, 363–385. 10.1146/annurev-vision-030320-041306.

49. Roe, A.W., Chelazzi, L., Connor, C.E., Conway, B.R., Fujita, I., Gallant, J.L., Lu, H., and Vanduffel, W. (2012). Toward a Unified Theory of Visual Area V4. Neuron. 74, 12–29. 10.1016/j.neuron.2012.03.011.

50. Fecteau, J.H., and Munoz, D.P. (2006). Salience, relevance, and firing: a priority map for target selection. Trends in Cognitive Sciences 10, 382–390. 10.1016/j.tics.2006.06.011.

51. Kim, B., and Basso, M.A. (2008). Saccade Target Selection in the Superior Colliculus: A Signal Detection Theory Approach. J Neurosci 28, 2991–3007. 10.1523/JNEUROSCI.5424-07.2008.

52. Boye, S.E., Alexander, J.J., Witherspoon, C.D., Boye, S.L., Peterson, J.J., Clark, M.E., Sandefer, K.J., Girkin, C.A., Hauswirth, W.W., and Gamlin, P.D. (2016). Highly Efficient Delivery of Adeno-Associated Viral Vectors to the Primate Retina. Hum Gene Ther 27, 580–597. 10.1089/hum.2016.085.

53. Cang, J., and Feldheim, D.A. (2013). Developmental Mechanisms of Topographic Map Formation and Alignment. Annu. Rev. Neurosci. 36, 51–77. 10.1146/annurev-neuro-062012-170341.

54. Lemke, G., and Reber, M. (2005). RETINOTECTAL MAPPING: New Insights from Molecular Genetics. Annu. Rev. Cell Dev. Biol. 21, 551–580. 10.1146/annurev.cellbio.20.022403.093702.

55. Weiskrantz, L. (2004). Roots of blindsight. Prog Brain Res 144, 229–241. 10.1016/s0079-6123(03)14416-0.

56. Takaura, K., Yoshida, M., and Isa, T. (2011). Neural Substrate of Spatial Memory in the Superior Colliculus after Damage to the Primary Visual Cortex. J Neurosci 31, 4233–4241. 10.1523/JNEUROSCI.5143-10.2011.

57. Takakuwa, N., Kato, R., Redgrave, P., and Isa, T. (2017). Emergence of visually-evoked reward expectation signals in dopamine neurons via the superior colliculus in V1 lesioned monkeys. Elife 6, e24459. 10.7554/eLife.24459.

58. Bridge, H., Leopold, D.A., and Bourne, J.A. (2016). Adaptive Pulvinar Circuitry Supports Visual Cognition. Trends in Cognitive Sciences 20, 146–157. 10.1016/j.tics.2015.10.003.

59. Eastman, K.M., and Huk, A.C. (2012). PLDAPS: A hardware architecture and software toolbox for neurophysiology requiring complex visual stimuli and online behavioral control. Frontiers in Neuroinformatics 6, 1. 10.3389/fninf.2012.00001.

60. Malpeli, J.G., and Baker, F.H. (1975). The representation of the visual field in the lateral geniculate nucleus of *Macaca mulatta*. J of Comparative Neurology 161, 569–594. 10.1002/cne.901610407.

61. Fisher, R.A. (1932). Statistical Methods for Research Workers 4th ed.

62. Bohlen, M.O., Rudzite, A.M., Daw, T.B., Kuczewski, G.M., Spiro, E., Hammond, C., Rogers, D.R., Gallego-Ortega, A., Manookin, M.B., Roy, S., et al. (2026). Projection targeting with phototagging to study the structure and function of retinal ganglion cells. Cell Reports Methods 6. 10.1016/j.crmeth.2026.101308.

63. Pachitariu, M., Steinmetz, N., Kadir, S., Carandini, M., and Harris, K. (2016). Fast and accurate spike sorting of high-channel count probes with KiloSort. Advances in neural information processing systems, 4448.

64. Kelly, R.C., Smith, M.A., Samonds, J.M., Kohn, A., Bonds, A.B., Movshon, J.A., and Lee, T.S. (2007). Comparison of recordings from microelectrode arrays and single electrodes in the visual cortex. The Journal of Neuroscience 27, 261–264. 10.1523/JNEUROSCI.4906-06.2007.

